# Myofascial Trigger Point (MTrP) Size and Elasticity Properties Can Be Used to Differentiate Active and Latent MTrPs in Lower Back Skeletal Muscle

**DOI:** 10.1101/2023.10.14.562354

**Authors:** P. Tsai, J. Edison, C. Wang, J. Sefton, K. Manning, M.W. Gramlich

## Abstract

Myofascial Trigger Points (MTrPs) are localized contraction knots that develop after muscle overuse or an acute trauma. Significant work has been done to understand, diagnose, and treat MTrPs in order to improve patients suffering from their effects. However, effective non-invasive diagnostic tools are still a missing gap in both understanding and treating MTrPs. Effective treatments for patients suffering from MTrP mediated pain require a means to measure MTrP properties quantitatively and diagnostically both prior to and during intervention. Further, quantitative measurements of MTrPs are often limited by the availability of equipment and training. Here we develop ultrasound (US) based diagnostic metrics that clinicians can use during patient diagnosis and treatment. We highlight the advantages and limitations of previous US-based approaches that utilize elasticity theory. We show how US-based measurements can distinguish *Active* from *Latent* MTrPs. We demonstrate that Active MTrPs tend to be smaller, stiffer, and deeper in the muscle tissue. We provide evidence that more than one MTrP within a single US-image field increases the stiffness of neighboring MTrPs. Finally, we highlight a combination of metrics (depth, thickness, and stiffness) that can be used to assess individual MTrPs in combination with standard clinical assessments.

## Introduction

It is estimated that 30%-85% of patients visiting primary care or pain clinics suffer from myofascial pain syndrome (MPS), a painful condition that affect muscles and fascia.^1,2^ Small nodules of tight tissue, called myofascial ^3,4^trigger points (MTrPs), can be found in the affected muscle tissues.^1,2^ Diagnosis of MPS is primarily based on patients” report and physical examination, such as palpation. This presents a serious problem for proper diagnosis since effective palpation requires clinicians to possess vital skills and experience. Like most chronic pain conditions, patients may first be seen by primary physicians. Unfortunately, most primary physicians lack the skill and experience to identify MPS and intervene. In addition, most patients may be reluctant to report or dismiss their suffering until it flares up and requires urgent medical attention. Therefore, similar to other chronic pain conditions, MPS patients may potentially be undertreated.^5^ However, until the field can be advanced to provide proper MPS diagnosis, we cannot be certain about the prevalence of MPS and its consequences. This lack of advancement also prevents the development of training protocols for providers to identify and treat MPS and teaching materials for patients to be made aware of this chronic pain condition and treatment options. Therefore, the overarching goal of this project was to identify an objective measurement protocol for MPS diagnosis and treatment outcomes evaluation. In this study, we used MPS of the low back as a study model since it is one of the most affected body regions.^3,4^

It has been hypothesized that poor posture, muscle overuse and/or physical injury lead to muscle overload. This creates a series of responses, including an increase of local acetylcholine (ACh) levels that may create pain and discomfort resulting in abnormal ACh release. ACh release is associated with ischemia and hypoxia in muscle tissue, disrupted mitochondrial activities, and the release of sensitizing substances.^6^ Sustained muscle contraction and continuous release of these molecules often cause nociception and pain reaction.^6,7^ MTrPs form as a local contraction in a small number of muscle fibers in a larger muscle bundle or muscle mass, which in turn can pull on tendons and ligaments associated with the muscle. All of these factors reduce muscle strength and alter the elastic capacity of the affected muscle.^6,7^

The elastic properties of MTrPs have been established as biomarkers because they can provide insights into MTrP formation and effectiveness of intervention treatments. Fundamental elasticity theory^8^ has been an effective framework to understand and quantify MTrP properties. MTrPs are composed of surrounding elastic muscle tissue which have established elasticity properties.^9^ MTrPs function as defects in the local muscle structure,^10,11^ equivalent to point defects in a lattice structure.^8,12^ This has proven to be a useful framework. MTrPs have been observed in vivo imaging to have fewer elastic properties than the surrounding tissue.^10^ Previous studies have assumed the MTrP defects and surrounding tissues are both composed of homogenous material and structure.^10^ This may not be accurate, thus limiting the application of the theory. Further, elasticity theory states the local defects also alter the elasticity properties of the surrounding environment,^12^ implying that developing and worsening MTrPs may be associated with a change of elasticity in the affected tissues. Therefore, identifying biomarkers that can be used to better quantify levels of elasticity in both MTrPs and surrounding muscle tissues are essential to assist in diagnosing MTrP(s).

Recent developments indicate that structural imaging methods are effective tools to identify MTrPs. Ultrasound (US) is one of the most promising tools. US can be used to differentiate between MTrPs and the surrounding normal tissues,^11^ and to map the elastic properties of soft tissues.^10^ US imagery indicates MTrPs are much stiffer than normal tissue in terms of tissue strain.^11,13–15^ However, recent research using vibration sonoelastography has failed to demonstrate the sensitivity to distinguish between *Active* and *Latent* MTrPs.^16^ The data showed that active and latent MTrPs have a similar degree of stiffness.^5^ This presents a major challenge for early treatment for patients with latent MTrPs. Latent MTrPs are often not associated with spontaneous pain report,^41^ which may prevent patients from seeking medical attention. Delayed detection and treatment of latent MTrPs could lead to worsening of MPS and further development of chronicity.

MTrP locations also affect the ability to diagnose and develop effective intervention techniques. MTrPs are often observed around the head, neck,^11^ shoulder, back.^10^ and extremities. MTrPs in these locations have different environments (composed of muscle, fascia and connective tissue) that can in turn affect their properties: size, elastic properties, amounts of blood flow in the surrounding tissue, and even the ability to be effectively imaged. Effective treatments for MTrPs will change depending upon location due to how the local muscles are being used.

Gaps in our understanding remain despite broadly established knowledge of the pathophysiology of MTrPs and the application of US for their diagnosis and treatment. Previous work exploring elasticitic properties of muscle explored *in vitro* muscle tissue revealed vast differences in native elasticity.^9,17^ Additionally, in vivo measurements of MTrP within muscle tissues were based on assumptions involving elastic properties of MTrPs and muscles tissue, without direct measurements.^10,11^ Further, the application of force-based intervention treatment, such either static force^10^ or vibrational waves,^11,18^ did not take into account how changes in the local tissue environment alter results. Thus, a comprehensive US and force-based approach that can be utilized by clinicians is still a major gap.

In the present study we quantitatively differentiate the elastic properties of MTrPs in lower back muscles using a US and static force-based approach. We first establish that all MTrPs respond to a localized static compressive force by establishing a stiffness parameter that is directly measurable using US imaging. We then categorically distinguish *Active* from *Latent* MTrPs, as well as potential atypical patient responses. Next, we utilize the stiffness metric to show that the MTrP response to applied force depends upon: (i) MTrP depth in muscle tissue, (ii) MTrP size relative to muscle tissue, and (iii) MTrP distance from applied force. We show that the number of MTrPs affect the elasticity properties of each individual MTrP. Finally, we present how stiffness and corrected strain parameters can distinguish the difference between *Active* and *Latent* MTrPs in a clinically relevant diagnosis environment.

## Results

### 1. MTrP Identification and Quantification

We first established the parameters used to measure and quantify MTrP response to pressure (Fig. 1). MTrPs were identified using an established palpation approach (see methods). We defined the longitudinal direction along the muscle fiber, and transverse as perpendicular to that direction. We then quantified the transverse local muscle tissue size (Tissue Size, **Fig. 1 A**) to determine the location of the MTrP within the muscle. We quantified the depth of the MTrP (MTrP depth, **Fig. 1 A**) relative to the top of the muscle tissue without applied pressure; this provides a standard approach from which to measure changes due to applied pressure. Finally, we quantified the transverse height of the MTrP (height, **Fig. 1 A**) at its center. These combined measurements provide sufficient metrics from ultrasound image analysis for use in MTrP diagnostic analysis.

**Figure 1:**
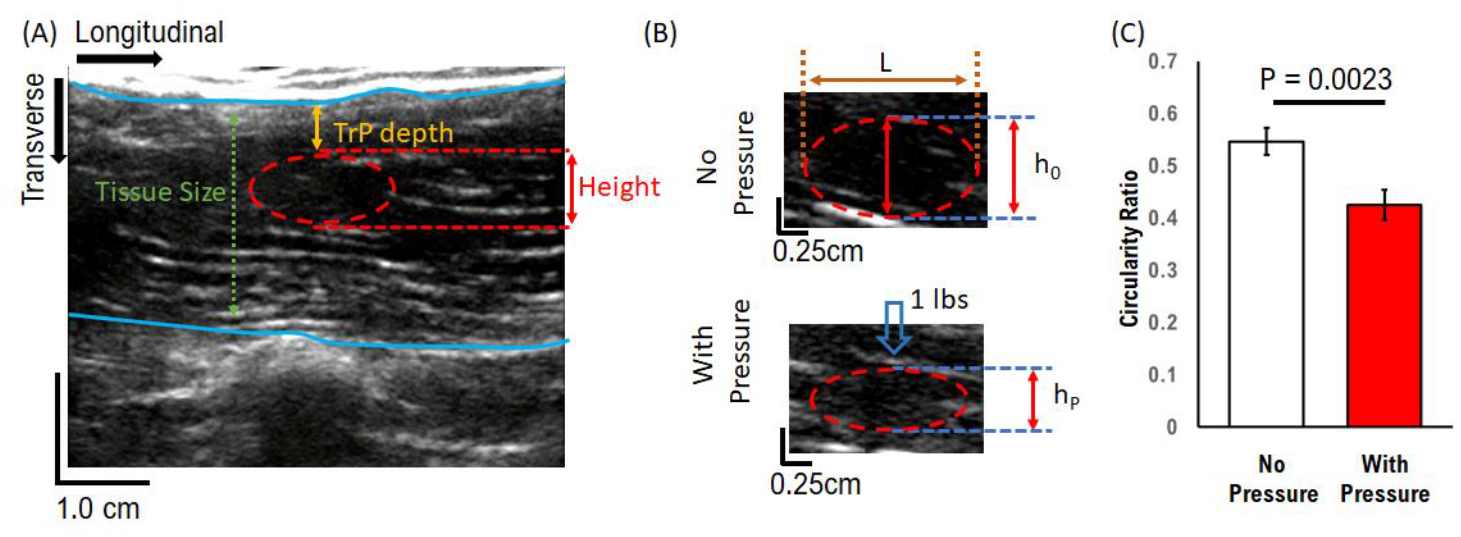
Experimental MTrP measurement resolution and Analysis: (A) Example MTrP measurement using ultrasound. MTrP point measurements are identified relative to the muscle tissue size. (B) MTrP compression resolution is measured using transverse height of a single MTrP and its corresponding longitudinal length (L). Measured height (h_0_) decreases under direct application of 1 lbp pressure (h_p_). (C) Aggregate circularity ratio (h/L) decreases with pressure. ** = P < 0.01, 2-tailed t-Test; Mean +/-SEM; N = 65 both conditions

To confirm that our approach was sufficient to distinguish changes in MTrPs under pressure, we used the circularity of the MTrPs before and after applied pressure. Typically, MTrPs are elliptical in shape (Top Panel, **Fig. 1B**), with longitudinal length (L) approximately twice the transverse height (h_0_) (Circularity 0.54 +/-0.027, **Fig. 1 C**) that is aligned with the direction of muscle contraction. Under an applied static local pressure of 1 lbs. transverse to the muscle, MTrP transverse height significantly reduced in circularity ratio by ∼22% (0.42 +/-0.03; P = 0.0023, two-tailed t-Test). We note here that this difference is based on MTrP response to an applied force directly above the MTrP location, and we will show later that this ratio decreases with increasing longitudinal distance between the location of the applied force and the MTrP. This measured reduction shows that transverse height measurement is a basis for quantifying changes in MTrPs under applied pressure using ultrasound.

In the present study, we used a single measure of MTrP response to apply pressure in order to quantify the physical properties of MTrPs. Typically, MTrP analysis uses traditional Young”s Modulus calculations from elasticity theory to determine elastic properties, which is defined as:

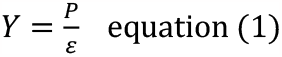

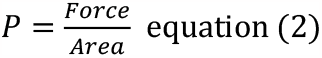

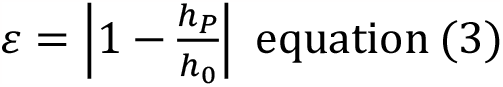

where h_p_ is height after pressure (hp, Fig. 1B) and h_0_ the height before pressure (h_0_, Fig. 1B).

This approach assumes applied pressure is uniform at the MTrP regardless of the properties of the muscle tissue and MTrP position, which we will show is not always accurate. Instead, in this study we used the height ratio as our metric, which we call *stiffness*, and define simply as the height of the MTrP with pressure divided by the height of the MTrP without pressure. This *stiffness* parameter does not require any assumptions about how the MTrP responds to applied force. We interpret this metric such that a ratio of 1 implies that the MTrP does not respond to applied force and is completely stiff, while a ratio approaching 0 implies that the MTrP compresses under applied force and thus more elastic.

### 2. Patient MTrP categories of Response

We tracked patient pain and twitching responses in addition to the US measurements. The existence of an MTrP does not necessarily guarantee that an individual will report spontaneous pain. Further, under locally applied pressure affected muscles may or may not twitch in the presence of a nodule confirmed by palpation and ultrasound procedure. We collected 75 ultrasound images and then designated four distinct categories based on Travell and Simons criteria (Table 1).^6^ One outlier was identified as the image which included three MTrPs, and was removed from data analysis.

**Table 1:** Pain and twitching categories.

| Category | Report of Spontaneous Pain | Twitching Observed | N<sub>single</sub> | N<sub>double</sub> |
| --- | --- | --- | --- | --- |
| 1 (active) | Yes | Yes | 16 | 3 |
| 2 (latent) | No | Yes | 8 | 0 |
| 3 (Atypical active) | Yes | No | 15 | 2 |
| 4 (Atypical latent) | No | No | 20 | 9 |

We compared the MTrP *stiffness* parameter across categories to determine if it is capable of discerning differences in MTrPs using two approaches. First, we compared MTrPs from all patients with reported pain (Category 1 & 3) and from patients reporting no pain (Category 2 & 4). Second, we compared MTrPs from patients with measured twitching (Categories 1 & 2) and from patients that have no measured twitching (Categories 3 & 4). Finally, we compared all four categories separately. This hierarchical approach was used to establish the sensitivity of US and *stiffness* as diagnostic tools for clinicians.

### 3. MTrP Response to Applied Static Force Depends on MTrP Depth in Tissue

We characterized the response of MTrPs to applied static force as a function of depth within the tissue, which has several clinical implications. First, palpation procedure applies force directly on the muscle surface. Potentially MTrPs near the surface may experience more force leading to a twitching response while MTrPs that lie deep within the muscle experience less force and may not produce a twitching response. The effectiveness of any static force-dependent treatment should also be affected by the location of MTrPs in the tissue. For example, force used to treat MTrPs is usually applied to the tissue surface (**Fig. 2A**). Surface applied force must then translate through the muscle to the MTrP within the tissue. Previous studies assumed that the force at the MTrP was equal to the applied force at the surface;^10,16,19^ however, elasticity theory shows that an applied force at a localized point on the surface of any material will begin to decrease as a function of depth following Saint-Venant”s principle.^20^ This would mean that force experienced locally within the muscle tissue will be less than the applied on the surface of muscle tissue, and would follow a quadratic relationship.

**Figure 2:**
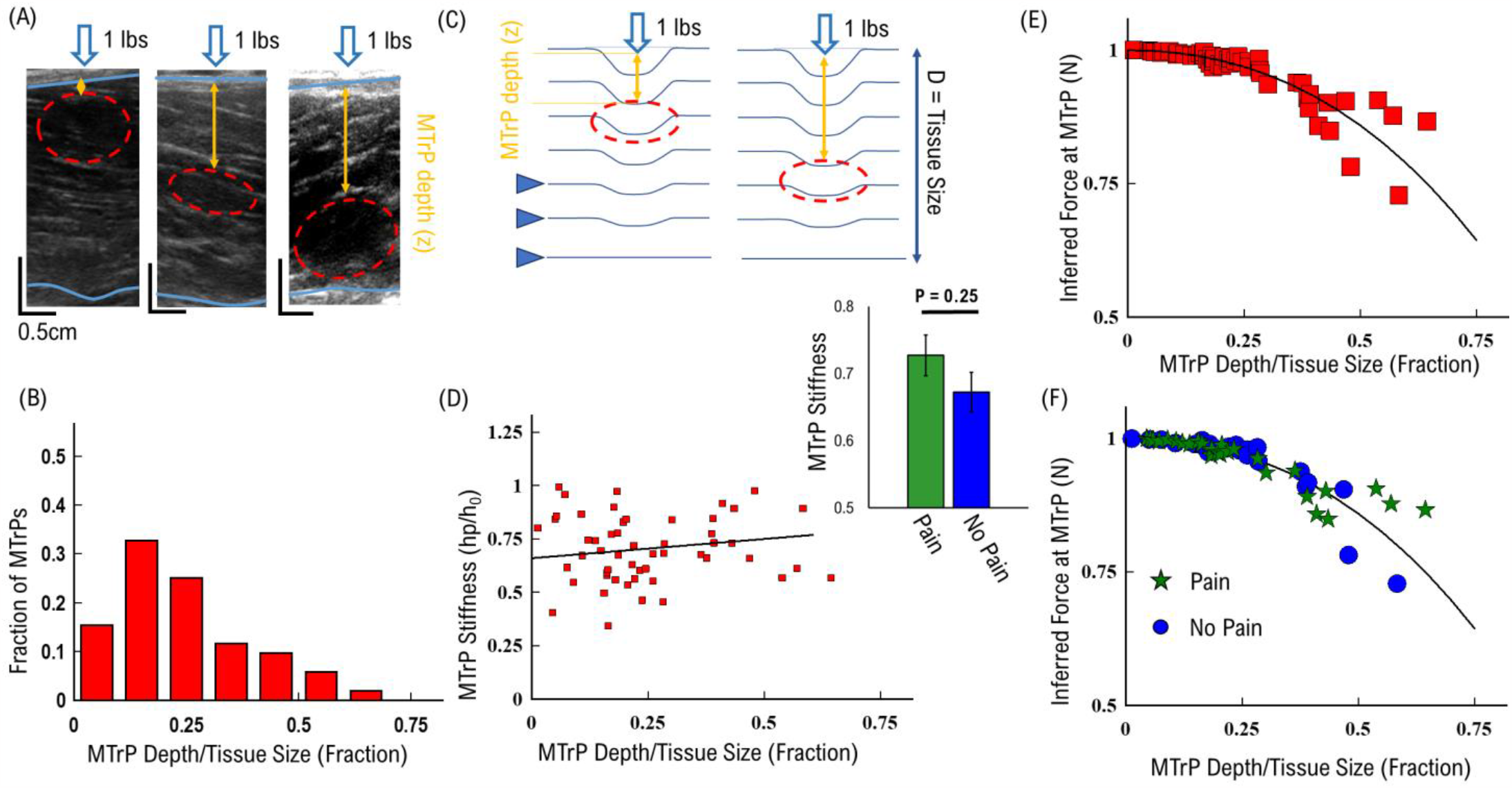
Single MTrP Response. (A) Model representation of decreasing deformation with depth. Local force experienced by the MTrP decreases quadratically with depth. (B) Distribution of MTrPs versus depth in tissue (C) Cartoon Model of MTrP compression dependent upon depth in tissue (D) MTrP stiffness increases linearly with depth in tissue. Inset shows average MTrP stiffness separated by Pain (Green) and No-Pain (Blue). (E) Inferred Effective for at MTrP as a function of depth in tissue. (F) Inferred Effective for at MTrP as a function of depth in tissue separated by Pain (Green Stars) and No-Pain (Blue Circles).

We established the depth of MTrPs within the muscle tissue in order to distinguish how they respond to applied force. We observed that MTrPs can exist at all depths within the muscle tissue (**Fig. 2A**). However, the majority of MTrPs (∼70%) occur near the surface of the muscle (Top 25% of muscle depth, **Fig. 2B**), which corresponds to the surface of the body. This suggests that the majority of MTrPs will not experience a significant reduction in applied force as previously assumed. Alternatively, a significant minority of MTrPs (∼30%) exist deeper into the muscle tissue (Bottom 75% of depth, **Fig. 2B**) and would thus experience a decreasing effective force with increasing depth.^20^

In order to determine if MTrP response to static force follows Saint-Venant”s principle, we modeled the muscle tissue as a stacked layer of equally elastic material (**Fig. 2C**). Applied static force at the surface deforms each muscle fiber layer which is then translated to a deformation force on the MTrP (Triangles indicating layers represented by deformed lines, **Fig. 2C**). This layered model also assumes that force is translated exclusively along the z-axis into the muscle, and only a fraction of force is translated between each muscle fiber layer, which we represent as decreasing layer deformations with increasing depth (Solid Lines, **Fig. 2C**). This model then predicts that the force at any given muscle layer decreases with depth and corresponds with a decrease in MTrP stiffness as a function of depth for an applied static surface force.

To test this response vs. depth relationship prediction, we measured MTrP stiffness (h_p_/h_0_, **Fig. 1B**) as a function of depth in the muscle (**Fig. 2D**). We observed that the majority of MTrPs near the surface have a mean stiffness of ∼0.67, and MTrPs exhibited an *apparent* slight correlation of increasing stiffness with depth in the tissue (Solid line, **Fig. 2 C**), with a rate of 0.2 per fraction of depth in tissue (R^2^ 0.08). The force at the surface is static. The apparent increasing stiffness is likely a consequence of decreasing force experienced at the MTrP within the muscle tissue.

We then use MTrP observed stiffness to determine an inferred depth-dependent local force to further support the depth-dependent MTrPs results. If we assume that the average change in MTrP stiffness corresponds to a decrease in the local effective force experienced at the MTrP, then we can use the measured stiffness results to estimate what the effective force on the MTrP would be as a function of depth. To perform this inference, we re-scaled the stiffness values by their effective depth following a quadratic relationship:

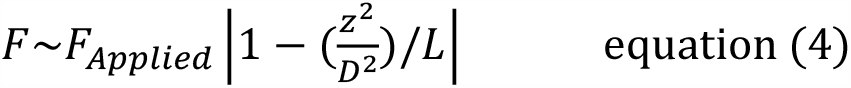

where z is the depth of the MTrP in the tissue and D is the total diameter of the muscle tissue, L is the characteristic decay length over which the force will dissipate. If the data follows Saint-Venant”s principle, then the observed stiffness values should all collapse onto the same quadratic curve above.

We found that the majority of MTrPs collapse onto the expected quadratic relationship with depth according to Saint-Venant”s principle (**Fig. 2E**). To apply this relationship, we multiplied each stiffness value (Red squares, **Fig. 2D**) by its normalized depth, squared the result, and subtracted from one. First, the average relationship of stiffness with depth rescales quadratically as the inferred force (solid line, **Fig. 2E**). Second, the rate of change in stiffness with depth (0.2, **Fig. 2D**), now represents the characteristic decay length (L) over which force dissipates within muscle tissue. Third, the majority of MTrPs re-scale onto the average inferred force (Red squares, **Fig. 2E**), which corresponds with the majority of MTrPs near the surface of the muscle tissue (**Fig. 2B**). This result suggests that the majority of MTrPs near the surface (<40%, **Fig. 2E**) follow the quadratic relationship.

Rescaled effective MTrPs *stiffness* values began to deviate away from the re-scaled effective force (solid line, **Fig. 2E**) with depth greater than 40% of the muscle tissue. To determine if the deviation was due to elasticity differences in category, we separated the results by the Pain/No-Pain groups (Table 1). First, we measured overall stiffness (independent of depth) separated by Pain/No-Pain categories (Inset, **Fig. 2D**). We found that on average patients with pain have MTrPs with a slightly higher stiffness (0.73 +/-0.03) compared to patients with no-pain (0.67 +/-0.03). To determine if these average differences are reflected in the depth-dependent results, we re-scaled the stiffness values as above (eqn. 4) and separated by Pain/No-Pain groups. We observed that the deviation from the average inferred force was different for patients with pain and patients with no-pain (**Fig. 2F**). Beyond a depth of 30%, MTrPs from patients with reported pain remained above the average inferred force (Green Stars, **Fig. 2F**), while MTrPs from patients with no-pain reported remained below the average inferred force (Blue circles, **Fig. 2F**). We assume that there is no difference in the local force for either groups, and thus this result suggests that patients that report pain have MTrPs with a higher stiffness than MTrPs from patients with no-pain reported, for the same force and depth confirming the finding of Fig. 2D.

Taken together, these results suggest that the depth of any MTrP must be considered as a parameter in determining any force-dependent intervention. Further, the depth-dependent results (**Fig. 2 D, F**) support the hypothesis that MTrPs have different elastic properties, and that their elastic properties mediate whether patients report pain or no-pain.

### 3. MTrP Elasticity is Dependent on MTrP Size and Patient Pain Report

We determined if MTrPs stiffness response to applied force depends upon size of the MTrP. Ultrasound results show that MTrPs exhibit different sizes and orientations within muscle tissue (**Fig. 3A**). Elastic material is typically assumed to be homogenous and isotropic, which would result in a compression response independent of size.^8,11^ This has been an implicit assumption in the context of quantifying the elastic properties of MTrPs. However, the pathophysiology of MTrP formation is hypothesized to result from a low pH, accumulation of Ca^2+^, recruitment of motor units, and dysfunctional actin/myosin cross-bridging.^21,22^ The consequence of this complex distribution of tissue and motor unit dysfunction would thus lead to the possibility of a heterogeneous structure that corresponds with MTrP size. This potential heterogenous combination of materials would result in elastic properties that change with size. Further, no previous study has directly explored MTrP elasticity as a function of size, which would have significant implications on designing diagnostic tools for MPS.

**Figure 3:**
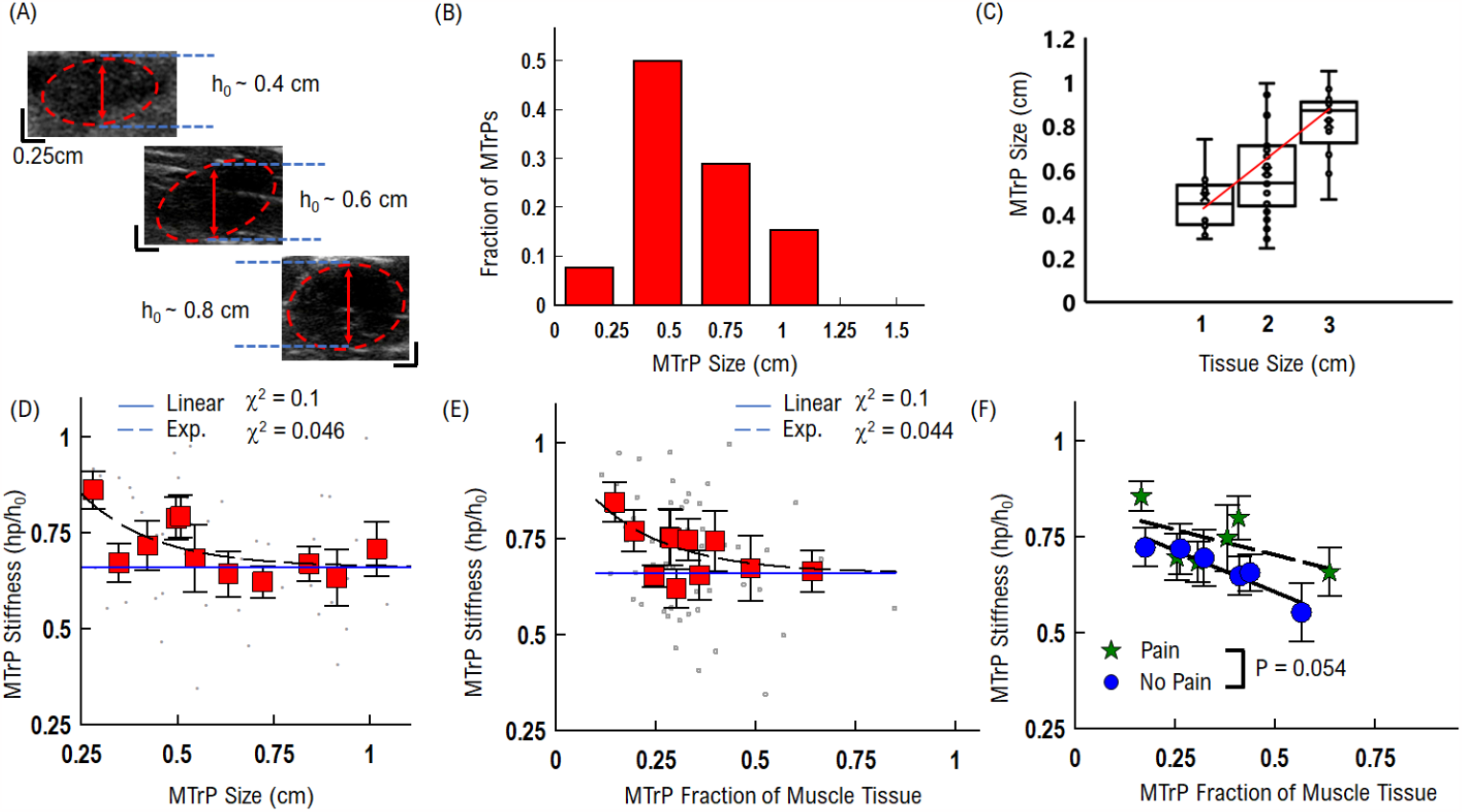
Single MTrP Stiffness Response Correlates with MTrP Size. (A) Example MTrP compression measured versus original MTrP thickness (B) Distribution of MTrP sizes. Average MTrP size is 0.6 cm. (C) MTrP size correlates with muscle thickness (D) MTrP stiffness slightly decreases with increasing thickness. Grey points show raw data, and Red squares are equal number averaged data. (E)MTrP stiffness slightly decreases with increasing Fraction of Muscle Tissue. Grey points show raw data, and Red squares are equal number averaged data (F) MTrP as a function of muscle fraction separated by Pain (Green Stars) and No-pain (Blue Circles). Data for each group have been separated into equal number (N = 6 MTrPs) bins. On average, Active MTrPs are more stiff than Latent MTrPs for the same size and fraction of muscle tissue. Statistical Comparison from pair-wise repeated measures t-Test.

We characterized MTrP size and correlation with muscle tissue size in order to establish how MTrP size distribution varies within the muscle. MTrPs are composed of muscle tissue and thus would be constrained by the total amount of muscle available.^21,22^ We observed that the average MTrP size was predominantly 0.6 +/-0.03 cm (**Fig. 3B**). We then observed that this average MTrP size depended upon the muscle tissue size, where the MTrP size increased linearly with muscle tissue size (0.2 cm increase in MTrP size per every cm increase of muscle tissue size, **Fig. 3C**). Further, the fraction of muscle tissue taken up by MTrPs is a significant amount for smaller muscles (40% for 1 cm thick muscle), but quickly decreases for larger muscles (25-30% for 3 cm thick muscles). These results suggest that MTrP stiffness and effects on surrounding muscle tissue will depend upon the MTrP size and fraction of muscle tissue it encompasses.

To understand if and how MTrP size affects elasticity measurements, we compared MTrP stiffness as a function of MTrP size and fraction of muscle tissue. We quantified MTrP stiffness with respect to its size independent of the surrounding tissue and found that stiffness decreased with increasing MTrP size (**Fig. 3D**). Average MTrP stiffness decreased exponentially with a rate of 0.25 cm (Dashed line, **Fig. 3D**), up to a thickness of 0.6 cm at which point MTrP stiffness remained constant (0.6, Solid Line, **Fig 3D**). The same relationship was observed with MTrP fraction of muscle size (**Fig. 3E**), where stiffness decreased exponentially (rate constant ∼0.2) with the increasing fraction of total muscle tissue it covered. These combined results that have the same relationship for MTrP size and muscle thickness suggest that MTrPs stiffness is dependent upon MTrP size but not dependent upon the fraction of muscle it covers. Further, the observation that MTrP stiffness is constant above 0.6 cm suggests that the MTrPs become more homogeneous in structure with increasing size. However, these results do not distinguish how the MTrP structure or function mediate size-dependent stiffness.

To understand the variance in the relationship between MTrP stiffness, size, and fraction of muscle tissue, we separated MTrP stiffness by Pain/No-pain categories (**Fig. 3F**). We observed that on average patients that reported pain have MTrP stiffness (Green Stars, **Fig. 3F**) that is larger (∼15-20%) than patients that reported no-pain MTrPs (Blue Circles, **Fig. 3F**) for the same fraction of muscle tissue. Further, both types of MTrPs stiffness decrease with increasing fraction of muscle tissue with a slightly slower rate for Pain (0.26, R^2^ = 0.56) compared to No-pain (0.43, R^2^ = 0.95).

These combined results show that MTrP elastic properties are dependent upon size and fraction of muscle tissue displaced. Further, these results show that patients that reported pain have MTrPs that are more stiff and less elastic than patients that reported no-pain MTrPs.

### 4. MTrP Response Decreases with Increasing Longitudinal Distance from Applied Force

Applied static force at any elastic material surface will decrease significantly with longitudinal distance from the location of the applied force.^8,23^ This means that any defect within the material will experience a different effective force depending upon its longitudinal distance from the location of the applied force. Consequently, along with depth and MTrP size, the longitudinal location of the MTrP relative to the applied force will affect the measured stiffness using US imaging (**Fig. 4 A**). If not taken into account, this difference will lead to incorrect measurements of MTrP stiffness, elasticity, and effectiveness of any force-based intervention approach. Thus, we next characterized the effective MTrP stiffness and effective force at the location of the MTrP as a function of longitudinal distances from the applied force.

**Figure 4:**
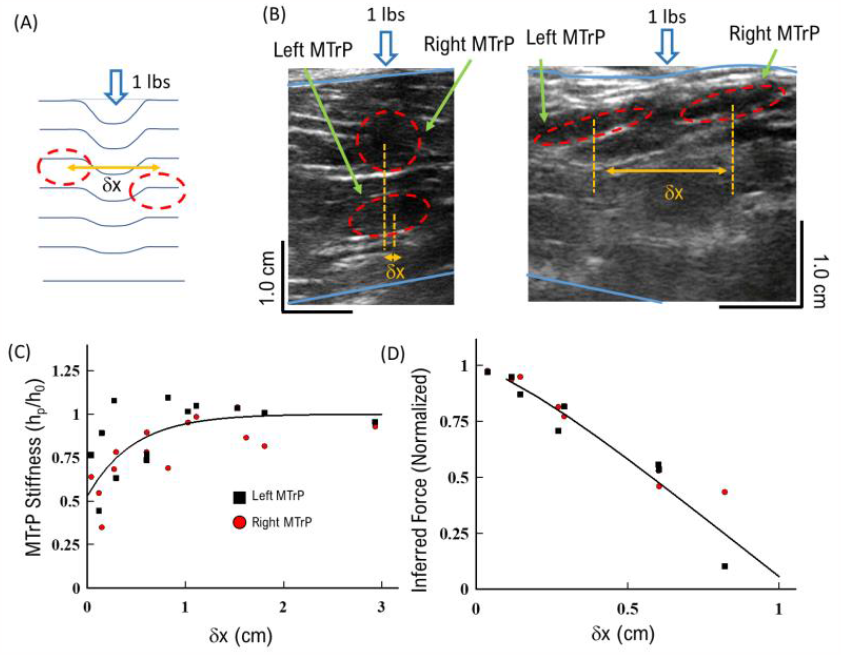
MTrP Stiffness Response Decreases with Increasing Longitudinal Distance from Applied Force: (A) Model of MTrP Response to applied static force as a function of distance from force (δx). (B) Example of two MTrPs directly beneath the applied force (Left Panel), or equal distance from applied force (Right Panel). Each MTrP is designated as *Left* or *Right*. (C) Stiffness of *Left* and *Right* MTrP versus longitudinal distance (δx) between the two MTrPs. Stiffness for both *Left* MTrP (Black Squares) and *Right* MTrP (Red Circles) decreases exponentially with distance between points (Solid Line). (D) Implied force on *Left* and *Right* MTrPs as a function of lateral distance from applied force

To determine the role of distance from applied force, we analyzed US data that included two MTrPs in a region (**Fig. 4 B, C**). This approach allowed us to distinguish the relationship between MTrP stiffness and applied force, because force is typically applied directly above a single MTrP; it may not be possible to apply force directly to all points for two or more MTrPs. Further, two MTrPs will allow us to distinguish any potential anisotropy in the elastic response of the muscle tissue as a function of direction. We applied a force at the center of each US image equidistant between two MTrPs for all patients imaged (**Fig. 4 B**). We quantified the distance between the two MTrP locations as the longitudinal distance from the applied force (δx, **Fig. 4 A, B**). We note that the distance of each MTrP is then half the longitudinal distance (δx/2).

We also distinguished the two MTrPs by direction in US images in order to discern any potential MTrP anisotropy in response to applied force. We defined the MTrP to the left of the force as *Left*, and the MTrP to the right of the force was defined as *Right*. If the two MTrPs laterally overlapped, then the bottom MTrP is defined as *Left* and the top MTrP is defined as *Right* (**Fig. 4B**). We note that we observed that the *Left* MTrP depth starts lower than the *Right* MTrP, per our definition (Left Panel, **Fig. 4B**), but *Left* MTrP depths decrease with increasing longitudinal distance to reach the same average depth as the Right MTrPs (Right Panel, **Fig. 4B**). This observation suggests that MTrPs in general have a preferential depth closer to the muscle surface, consistent with the single MTrP depth results (**Fig. 3B**).

We observed that both the *Left* and *Right* MTrP stiffness increased with distance from the applied force (**Fig. 4C**). First, both the *Left* and *Right* MTrP stiffness were lowest directly beneath the applied force (0.6 +/-0.07, **Fig. 4 C**), which is consistent with the single MTrP stiffness measurements (**Fig. 2D, 3D**). Second, the measured stiffness for both the *Left* and *Right* MTrPs quickly increases with increasing longitudinal distance to a maximum at approximately 1 cm away from the applied force (0.93 +/-0.057, **Fig. 4 C**). We then averaged the observed increase for both *Left* and *Right* MTrPs and observed the stiffness changed exponentially with increasing longitudinal distance (Solid line, **Fig. 4 C**). Since the applied static force is unchanged for all conditions, these results support our hypothesis that the effective force felt by each MTrP decreases with increasing longitudinal distance from the position of applied force (**Fig. 4 A**). We also note that both the *Left* and *Right* MTrPs have the same relationship between stiffness and distance from applied force, suggesting that there is no directional anisotropy in MTrP response and only the distance from applied force matters.

We applied the same quadratic re-scaling of MTrP following Saint-Venant”s principle (eqn. 1) to further support our hypothesis that differences in MTrP stiffness are due to changes in local effective force at the MTrP,.^23^ Here we assume that the effective force at the MTrP decreases quadratically with increasing longitudinal distance (δx, **Fig. 4 A, B**). We then scaled the observed stiffness for both Left and Right MTrPs and observed a significant decrease in the effective force with distance (δx, **Fig. 4 D**). The force experienced by any MTrP decreases 25% for every 0.25 cm away from the applied force, and all the applied force has been lost when the MTrP is greater than 1 cm away from the applied force. The consequence of this effective force result shows any intervention or diagnosis procedure that relies on applied pressure, such as palpation, will be limited by how closely the pressure is applied to the MTrP.

These combined stiffness and effective force results support the hypothesis that longitudinal distance from applied force significantly affects the observed stiffness, and distance must be considered when using US measurements and stiffness as diagnostic tools.

### 5. The Number of MTrPs in an US region Affects Measured Stiffness

We determined if there is a difference in how MTrPs respond to applied force for more than one MTrP in a given muscle region. Defects can change the elastic properties of the surrounding material, and thus influence other defects nearby.^8,12^ This occurs when one or more defects expands the surrounding material and alters the materials” ability to respond to applied static force. In the context of MTrPs, this would suggest that having more than one MTrP can influence the elastic properties of each compared to only having a single MTrP. Thus, any intervention or diagnostic procedure that relies on US measurements and applied force may report different effects depending on the numbers of MTrPs.

To determine if multiple MTrPs in a muscle alter the measured elastic properties of each individual MTrP, we compared both the strain response (ε, **eqn. 3**) and Young”s Modulus (Y, **eqn. 1**) of one MTrP to the two MTrP measurements (**Fig. 5 A,B**). Since depth and MTrP thickness would also influence the elasticity properties of the surrounding muscle tissue, we did not correct for either of these properties in order todetermine how much they influence resulting strain of more than one MTrP. However, we did correct the two MTrP data for distance from applied force (**Fig. 4**), in order to establish a native change at maximum force. Finally, we treated each *Left* and *Right* MTrP as separate defects so that 14 US images resulted in 28 unique MTrPs.

**Figure 5.**
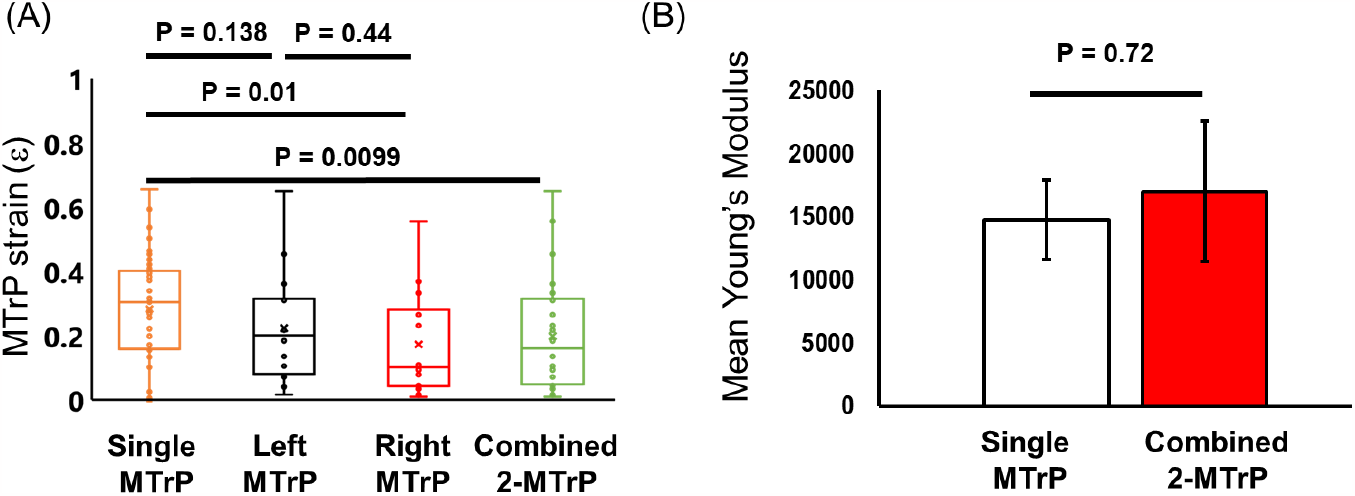
Comparison of one and two MTrPs within an US image region: (A) Comparison of strain for a single MTrP and two MTrP data. Single MTrPs are uncorrected, while Two MTrP data is corrected for longitudinal distance. (B) Calculated Young”s Modulus for single MTrP and two MTrPs. All data corrected for depth and distance from applied force. Means +/-SEM; Statistical tests are 2-tailed t-Test; N_single_ = 54; N_two_ = 28

We found that on average single MTrPs were more elastic than two MTrPs within a US image field (**Fig. 5 A,B**). We first observed that each *Left* and *Right* individual MTrP exhibited similar strains (*Left*: 0.22 +/-0.05; *Right*: 0.17 +/-0.04) to each other, supporting the result that there is not any significant anisotropy in MTrP response to applied force (**Fig. 4**). However, each *Left* and *Right* individual MTrP strains were both lower than single (0.29 +/-0.02) MTrP strain (**Fig. 5 A**). This difference is further highlighted when combining *Left* and *Right* MTrPs strain response (0.20 +/-0.03) compared to single MTrP strain, which showed that the average two MTrP strain response was significantly lower. This result suggests that each MTrP in the muscle influences the elastic properties of the surrounding muscle tissue and thereby affects the elastic properties of other MTrPs nearby.

We compared calculated Young”s Modulus for single and double observed MTrPs, but corrected for size and depth. This corrected MTrP analysis will provide a more accurate measurement of MTrP elasticity properties. First, we calculated the effective Tensile stress (See Methods) at the MTrP ∼3980 N/m^2^. Comparing both single and two MTrP data shows that the Young”s modulus (**Fig. 5B**) is slightly elevated for two MTrPs (Y = 17000 +/-5554) compared to a single MTrP (Y = 14300 +/-3125). Because of the variability in MTrP corrected strain response, this slight difference is not statistically significant. However, both single and two MTrP Youngs moduli are consistent with previously published results.^10^

These combined results suggest that more than one MTrP within a 1 cm muscle region influences the elastic properties of the surrounding muscle, and each other, resulting in greater stiffness.

### 6. Spontaneous Pain Report and Twitching as a Diagnostic of MTrP Properties

Finally, we focused on whether the properties of MTrPs can be used as diagnostic tools for both spontaneous pain report and measured twitching response. Because US MTrP elasticity measurements depend upon depth and size, it is possible that these same properties can be used as diagnostic tools for clinicians. For example, quickly identifying MTrP size and depth could be used to support other assessments of MTrP as either Active/Latent and the effectiveness of force-based treatments. We also note that we will only consider single MTrP results to distinguish MTrP properties with category.

We considered the native properties of MTrPs in the tissue for groups where twitching was observed, but without any response to applied force (**Fig. 6 A,B**). We observed a slight, but not significant, increase in MTrP thickness between patients that had a spontaneous pain report and twitching response (Group 1, *Active*) compared to patients that did not have a spontaneous pain report but twitching was observed (Group 2, *Latent*). In contrast, there was no significant difference in MTrP depth between *Active* and *Latent* groups (**Fig. 6B**). In the context of our observed relationship between MTrP stiffness and size, these results suggest that *Active* MTrPs will on average be slightly stiffer than *Latent* MTrPs, but not any deeper into the tissue.

**Figure 6:**
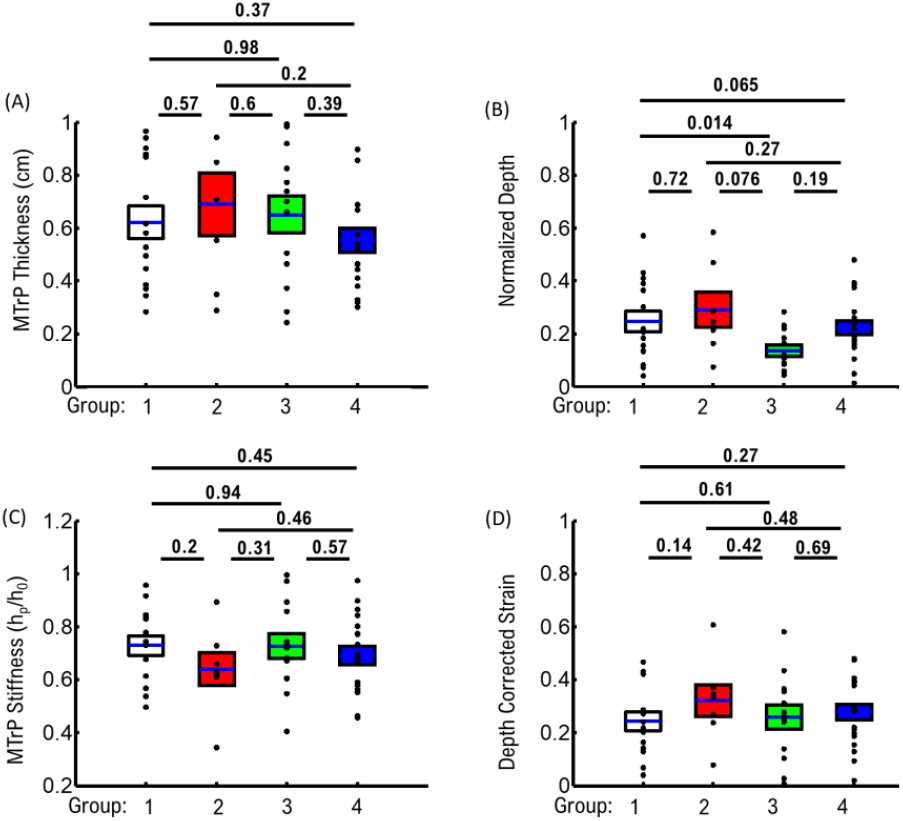
Comparison of MTrPs with Reported Pain and/or Twitching Group. (A) Comparison of MTrP thickness. (B) Comparison of MTrP Depth in tissue (C) Comparison of compression ratio for a single MTrP data. (D) Comparison of MTrP corrected strain All data corrected for depth and distance from applied force. All N-values given in table 1. Means +/-SEM; Statistical tests are 2-tailed t-Test;

When comparing groups exhibiting observed twitching (Groups 1+2) to groups where no twitching was observed (Groups 3+4), differences in MTrP properties were observed (**Fig. 6 A,B**). We observed that groups exhibiting twitching had slightly larger average MTrPs thickness (0.64 +/-0.04 cm), but not significant (P = 0.65, two-tailed t-Test), than groups that did not exhibit twitching (0.58 +/-0.03 cm). Further, we observed MTrPs from both twitching groups were deeper into the muscle tissue (0.26 +/-0.03 %) compared to MTrPs from no-twitching groups (0.19 +/-0.02 %) (P = 0.065, two-tailed t-Test). These combined results suggest that twitching response correlates with MTrPs that are slightly larger and deeper within the muscle tissue.

We considered the response of MTrPs to applied force as a diagnostic tool. We initially compared uncorrected stiffness between *Active* and *Latent* MTrPs to determine if MTrP response is depended on the native depth and thickness conditions (**Fig. 6C**). *Latent* MTrPs showed a slight, but not significant, reduction in *stiffness* (0.64 +/-0.06) compared to all other groups (Group 1 = 0.73 +/-0.04; Group 3 = 0.72 +/-0.04; Group 4 = 0.67 +/-0.04). However, when we compared strain corrected for depth (**Fig. 6D**), we observed that *Active* MTrPs showed a slight reduction (0.27 +/-0.03) compared to *Latent* MTrPs (0.36 +/-0.06). These results support the observation that *Active* MTrPs are less elastic than *Latent* MTrPs, and that depth-corrected strain is a useful diagnostic tool to distinguish the two different groups.

We distinguished changes for groups exhibiting twitching compared to no-twitching groups, to determine if force-dependent MTrP properties were dependent upon twitching. Un-corrected stiffness for twitching groups showed a larger stiffness response (0.72 +/-0.03) compared to no-twitching groups (0.68 +/-0.03), which is consistent with the average difference in depth between groups (**Fig. 6B**). The difference in stiffness also corresponds with a lower corrected strain (P = 0.22) observed for groups with twitching (0.27 +/-0.03) compared to groups without twitching (0.34 +/-0.03). These results suggest that stiffness and strain are potential diagnostic tools to distinguish MTrPs that result in twitching versus no-twitching.

The combined native MTrP properties (Thickness and Depth) and their respective elastic response to statice force (stiffness and strain) can be used as effective diagnostic tools to support clinical diagnosis and treatment.

## Conclusions

In the present study we utilized an US-based measurement approach to develop physical parameters to characterize MTrPs (**Fig. 1**). We established important limitations and considerations of MTrP properties such as depth in tissue (**Fig. 2**) and MTrP thickness (**Fig. 3**) when characterizing MTrP elasticity properties. We showed that Active MTrPs are more stiff and less elastic than Latent MTrPs (**Figs. 3, 6**). We showed that the effectiveness of any applied force intervention approach decreases when the MTrP is farther away from the location of applied force (**Fig. 4**). We showed that two MTrPs within 1 cm of each other alter the elastic properties of each MTrP (**Fig. 5**). Finally, we showed how native MTrP depth, thickness, and elasticity properties can be utilized as diagnostic tools for clinicians (**Fig. 6**).

## Discussion

It is important to put the limitations of US-based measured MTrP elasticity properties in context, in order for clinicians to utilize MTrP elasticity for diagnostic and treatment assessment purposes. The difference between *Active* and *Latent* stiffness is ∼9 +/-1% (**Fig. 2D**). While this difference can have significant biological effects on muscle function, the relatively small-scale difference means that the ability to distinguish any individual MTrP as either *Active* or *Latent* requires assessing multiple parameters simultaneously. From US imaging alone, we have identified three combined metrics (depth, thickness, stiffness) that can be utilized. Combined, these metrics show that *Active* MTrPs tend to be simultaneously deeper, smaller, and stiffer than *Latent* MTrPs. When assessing each individual MTrP, these three metrics can be directly measured during US imaging by a clinician without significant interpretation. However, while the present study shows how these metrics can be utilized as diagnostic tools, a more thorough protocol and parameter threshold should be established in future studies to help clinicians quickly measure and determine whether an MTrP is active or latent.

It is also important to highlight the limitation of using a single static applied force in the elastic characterization of MTrPs in the present study. Our main scope with this study was to develop and establish metrics for US-based MTrP analysis as well as limitations with the US-based approach. We focused on a single applied static force at the center of the US image (**Fig. 1**). This was sufficient to distinguish differences in Pain and No-Pain group MTrP elasticity properties as a function of depth (**Fig. 2**) and thickness (**Fig. 3**). However, elastic deformations can also have force-dependent differences due to heterogeneously different structures. We hypothesized in the current study that our observed relationship between MTrP size and stiffness (**Fig. 3**) is likely due to changes in the heterogeneous structure of MTrPs as a function of size. If we combine this hypothesis with our observation that *Active* MTrPs tend to be smaller than Latent MTrPs (**Fig. 6A**), then we would expect that *Active* and *Latent* MTrPs would also have different stiffness relationships with respect to applied force. This would represent another potential diagnostic tool to assess each individual MTrP and the effectiveness of any force-based intervention treatment. Future studies could explore this potential by measuring MTrP stiffness for different applied static surface forces.

## Methods

### Patient Recruitment

#### Sample

This project was part of a larger study which investigated characteristics of patients with MPS of the low back. This project only reported the difference among active, latent and atypical MTrPs at the local MTrP level. The original study was a cross-sectional descriptive study conducted between 8-30-2021 and 6-30-2022 after receiving approval from IRBs of Edward Via College of Osteopathic Medicine (VCOM) and Auburn University. Twenty-five participants were recruited from VCOM and Auburn University campuses using a snow ball sampling method. The study included participants who (1) were diagnosed with non-specific low back pain,

(2) English-speaking, and (3) ambulatory without a cane or walker. Participants were excluded if they had (1) major illness, such as cancer, (2) major surgery within 6 months, (3) major psychiatric disorder, such as bipolar disorder and depression, (4) cognitive impairment or (5) other painful conditions of the low back.

### Ultrasound Measurements

#### Identifying location, number and type of MTrPs

The participant was asked to lie in a prone position on an examination table. The participant was examined by an osteopathic doctor using a physical examination and palpation to determine the presence, site, and number of MTrPs on low back muscles between L1 and S1. The ultrasound procedure described later was used to confirm the finding of MTrPs.

The physician then randomly selected three MTrPs in each participant for further ultrasound evaluation procedure using the Sonosit Edge II system. Active MTrP site (Group 1) was defined as spontaneous local or referred pain on the examining day and a twitching response when palpating based on the definition of Travell and Simons^6^. A latent MTrP (Group 2) is similar to an active MTrP but no spontaneous local or referred pain on the examining day. However, when a nodule was identified by palpation and confirmed by US procedure but no twitching response, this study defined it as an atypical MTrP since it did not meet the definition proposed by Travell and Simons.^6^ This included sites that had “spontaneous pain but no twitching response when palpated” (Group 3), or no spontaneous pain and no twitching response when palpated” (Group 4), but tense nodules were readily visualized under ultrasound-guided examination.

Two images were taken for each MTrP site. One gray scale US image was taken without force and a second image was taken immediately after applying ∼4.5 N weight to the same site. Images were coded for follow-up analysis. We recorded coordinates and the length of the maximum vertical line of the MTrP muscle tissues with and without applied force. The measured length of maximum vertical lines in images with and without applied force was used to assess strain of muscle tissues. Stress was determined by dividing applied force by transducer contact area. The elastic modulus was determined by the ratio between stress and strain.

### Calculated Tensile Stress Due to Applied Force on the Ultrasound Head

Based on the reduction in measured MTrP compression with distance (**Fig. 4**), we corrected the total area over which our applied force occurred, which is less than the head size (See Methods); we assume that the surface area of the force applied on the muscle tissue occurs with a radius between 1.5 – 2 cm, given that measured MTrP compression did not occur beyond this distance.

### Exclusion Criteria for Data Analysis

MTrPs were excluded from analysis if their transverse height was greater in the presence of applied force than without applied force. This increase in height with applied force is likely a consequence of changing MTrP orientation and/or muscle orientation during US imaging.

### Statistical Analysis

Statistical comparisons between pressure/no-pressure (Fig. 1), and all MTrP groups (Fig. 2, 5, 6) were performed as pair-wise two-tailed t-Tests. Comparison between MTrP groups as a function of depth or thickness (Fig. 2, 3) were performed using repeated-measured t-Tests. Comparisons between MTrPs and equations describing MTrP stiffness as a function of depth/thickness were performed using χ^2^ analysis.

**Table 2:** Statistical P-values for Stiffness by group.

| Stiffness | 1 | 2 | 3 | 4 |
| --- | --- | --- | --- | --- |
| 1 | - | 0.2 | 0.94 | 0.45 |
| 2 |  | - | 0.31 | 0.46 |
| 3 |  |  | - | 0.57 |
| 4 |  |  |  | - |

**Table 3:**
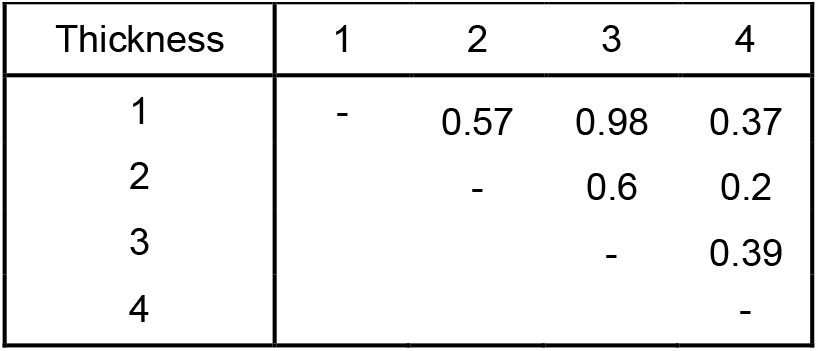
Statistical P-values for Thickness by group.

| Thickness | 1 | 2 | 3 | 4 |
| --- | --- | --- | --- | --- |
| 1 | - | 0.57 | 0.98 | 0.37 |
| 2 |  | - | 0.6 | 0.2 |
| 3 |  |  | - | 0.39 |
| 4 |  |  |  | - |

**Table 4:**
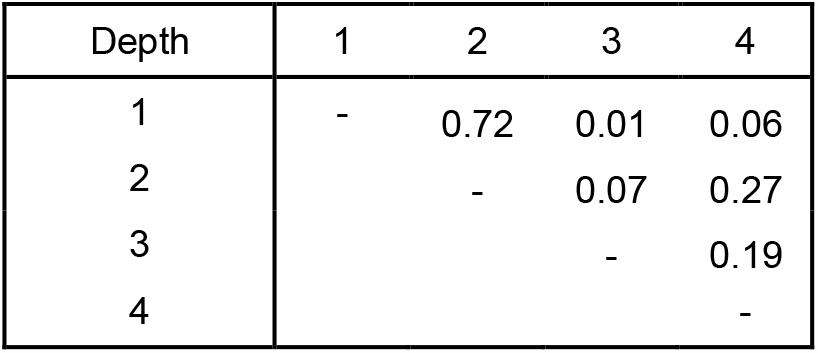
Statistical P-values for Depth by group.

**Table 5:**
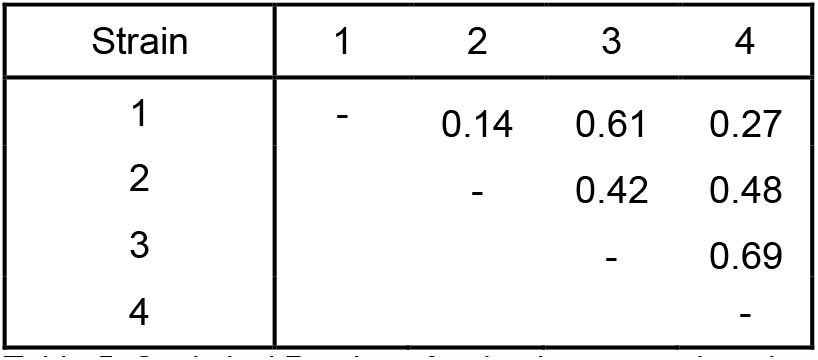
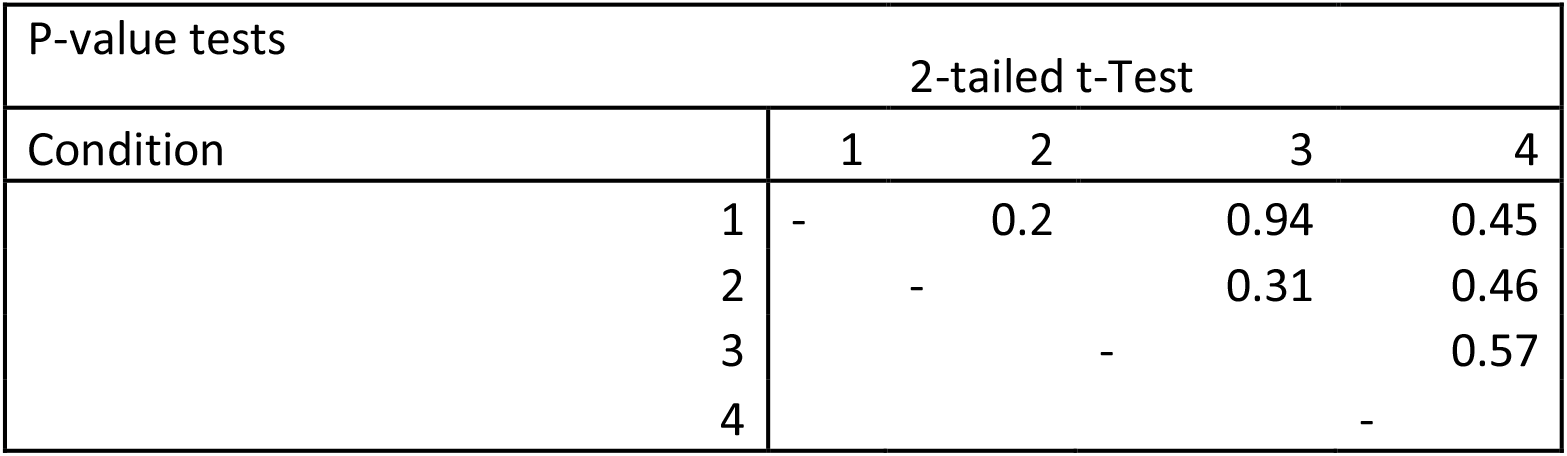
Statistical P-values for depth corrected strain by group.

## Author Contributions

PT, JE, CW, JT, and KM designed and carried out the study. PF, JE, KM, DB, CW, and MWG analyzed MTrP data. All authors contributed to the study design and writing of results from this study.

## Author Declarations

The authors do not declare any conflicts of interest in the present study.

## Acknowledgements

This study was supported by the Research Eureka Accelerator Program (REAP), Edward Via College of Osteopathic Medicine.

